# Quantitative spatial analysis of neuroligin-3 mRNA expression in the enteric nervous system reveals a potential role in neuronal-glial synapses and reduced expression in *Nlgn3^R451C^* mice

**DOI:** 10.1101/2022.05.02.490237

**Authors:** Madushani Herath, Ellie Cho, Ashley E Franks, Joel C Bornstein, Elisa L Hill-Yardin

## Abstract

Mutations in the Neuroligin-3 (Nlgn3) gene are implicated in autism spectrum disorder (ASD) and gastrointestinal (GI) dysfunction but its cellular expression in the GI tract remains to be characterised. Localisation of NLGN3 protein is challenging in intestinal tissue due to the lack of target-specific antibodies. Here, we combined RNAScope *in situ* hybridization for *Nlgn3* mRNA and immunofluorescence for markers of all enteric neurons, cholinergic submucosal neurons, non-cholinergic submucosal neurons, nitregic and calretinin-containing myenteric neurons as well as glial cells. We also developed a quantitative 3-dimensional image analysis method to measure *Nlgn3* mRNA cellular expression levels in enteric neurons and glia. We show that *Nlgn3* mRNA is expressed in most submucosal and myenteric neurons as well as in enteric glia. The R451C mutation reduces *Nlgn3* mRNA expression levels in cholinergic, nitrergic and calretinin enteric neuronal subpopulations but does not affect *Nlgn3* mRNA expression in VIPergic submucosal neurons. In summary, we show that the autism-associated R451C mutation in *Nlgn3* reduces *Nlgn3* mRNA expression in the mouse ENS. These findings could shed light on the pathophysiology of GI dysfunction in ASD.

## INTRODUCTION

Neuroligin-3 (NLGN3) is expressed in the central nervous system (CNS) and regulates brain function, but its cellular expression profile in the enteric nervous system has not been characterised. In the rat hippocampus, the NLGN3 adhesion molecule is expressed at the postsynaptic membrane of both excitatory and inhibitory synapses (Budreck and Scheiffele, 2007). It has been shown that the presence of NLGN3 is not restricted to neurons but also expressed in non-neuronal cells relevant to glioma (Venkatesh et al., 2015). Venkatesh and colleagues showed that neurally-secreted NLGN3 invades the microenvironment of tumours and induces NLGN3 expression in glioma cells to promote tumour growth in mice with high-grade glioma (Venkatesh et al., 2015). This evidence suggests that NLGN3 is not only involved in neuronal communication but also plays a role in neuron-glia interactions.

Mutations including the missense R451C point mutation as well as the ablation of the *NLGN3* gene are implicated in autism (Jamain et al., 2003). Patients expressing the *NLGN3* R451C mutation show GI dysfunction including diarrhea, faecal incontinence, post-meal regurgitation, oesophageal inflammation, chronic intestinal pain as well as delayed bladder and bowel control (Hosie et al., 2019). Modification of NLGN3 expression induces GI dysfunction in mice, as shown in both *Nlgn3* knockout (KO) and *Nlgn3^R451C^* mice (Leembruggen et al., 2019, Hosie et al., 2019). *Nlgn3* KO mice exhibit distended colons and faster colonic muscle contractions (Leembruggen et al., 2019). *Nlgn3^R451C^* mice display faster small intestinal transit and increased numbers of myenteric neurons alongside GABA_A_ receptor-mediated colonic dysmotility (Hosie et al., 2019) suggesting a role for NLGN3 in gut function.

Effects of the R451C mutation on *Nlgn3* mRNA and protein expression at the cellular level have been reported in the brain. In both HEK293 cell lines and mouse forebrain tissue, *Nlgn3* mRNA expression levels are unchanged by the *Nlgn3* R451C mutation but NLGN3 protein expression at the synaptic membrane is drastically reduced (Tabuchi et al., 2007, Comoletti et al., 2004). Although initial studies of NLGN3 expression in the GI tract have been reported in rodents and humans (Zhang et al., 2013, Hosie et al., 2019, Bohorquez et al., 2015), the subcellular distribution of NLGN3 expression in different cell types of the mouse enteric nervous system has not been profiled. Characterising *Nlgn3* expression in enteric cell subtypes is essential to better understand underlying mechanisms contributing to GI functional changes observed in patients and the *Nlgn3^R451C^* mouse model of autism.

Localizing the NLGN3 protein in the GI tract is challenging due to non-specific labelling of commercially available antibodies which commonly yield false-positive results (Leembruggen et al., unpublished). Therefore, an enhanced *in situ* hybridization technique (RNAScope) was pursued to label *Nlgn3* mRNA. RNAScope (ACD, 320850, USA) is designed to visualise individual RNA molecules using a novel probe design and amplification system to simultaneously magnify the signal and suppress background noise (Wang et al., 2012). Here we combined RNAScope and immunofluorescence with high-resolution microscopy and 3D image analysis software to characterise *Nlgn3* mRNA expression in the wild type and *Nlgn3^R451C^*mouse gut. Overall, we revealed that *Nlgn3* mRNA is expressed in the mouse enteric nervous system including glia, and that mice expressing the R451C mutation have reduced *Nlgn3* mRNA expression in enteric neurons in both the submucosal and myenteric plexus. Specifically, this mutation reduces *Nlgn3* mRNA expression levels in submucosal cholinergic, myenteric nitrergic and calretinin enteric neurons but does not affect *Nlgn3* mRNA expression in VIPergic submucosal neurons. The R451C mutation also reduces *Nlgn3* mRNA expression levels in myenteric but not submucosal glia. These findings provide further evidence that mutations in NLGN3 might play a role in gastrointestinal dysfunction in individuals with autism.

## METHODS

### Animals

B6;129-Nlgn3tm1Sud/J mice were obtained from The Jackson Laboratory (Tabuchi et al., 2007) and bred for over 10 generations on a pure C57BL/6 background at the Howard Florey Institute, Melbourne Australia. Mice were subsequently housed in the Biomedical Science Animal Facility at The University of Melbourne. *Nlgn3^R451C^* male mice and wild type (WT) mice (age 12-13 weeks) were killed by cervical dislocation in accordance with The University of Melbourne Animal Experimentation Ethics Committee (#1914843).

### Tissue preparation

*Nlgn3*^R451C^ male mice were killed and the abdomen of the animal was opened using coarse dissection scissors. A 2 cm segment of the distal ileum (1 cm proximal to the caecum) was isolated and immediately placed in oxygenated ice-cold physiological saline solution (composition in mM: 118 NaCl, 4.6 KCl, 2.5 CaCl_2_, 1.2 MgSO_4_, 1 NaH_2_PO_4_, 25 NaHCO_3_, 11 D-glucose). The intestinal content was gently flushed clean using a Pasteur pipette. Using small dissecting scissors, the distal ileal tissue sample was cut along the mesenteric border, stretched and pinned flat on a chilled silicon elastomer-lined dish (Sylgard 184, Dow Corning, Australia). The tissue was fixed in 4% formaldehyde for 24 h at 4°C. After fixation, the tissues were washed three times in 1M phosphate buffer saline (PBS) (composition in mM: 137 NaCl, 2.7 KCl, 8 Na_2_HPO_4_, 2 KH_2_PO_4_). The wholemount submucosal plexus preparation was isolated by peeling the mucosa-submucosal layer and then removing the epithelium using fine forceps (Fine Science Tools, Canada). Myenteric plexus preparations with the longitudinal muscle layer (LMMP) were obtained by microdissection to remove the circular muscle layer.

### Dual RNAScope *in situ* hybridization and immunofluorescence

RNAScope *in situ* hybridization in mouse ileal tissue was performed using the ACDBio multiplex RNAScope assay kit (ACD, 320850, USA) according to manufacturer’s instructions. Briefly, wholemount mouse ileal submucosal and LMMP preparations were pre-treated and hybridized with the RNAScope probe of interest. The hybridization signal was subsequently amplified via a sequence of amplifiers and fluorescently labelled probes.

### 2.3 Pre-treatment

Wholemount preparations of submucosal and LMMP preparations were permeabilized via pre-treatment with protease IV prior to RNAScope *in situ* hybridisation. The tissue samples were incubated in 30 μl of protease IV for 30 min at room temperature (RT) in a sealed humid chamber and washed twice (1 min in each) in 1M PBS.

### 2.4 RNAscope *in situ* hybridization

Following pre-treatment, ileal tissue preparations were incubated in 20 μl of *Nlgn3* RNAScope probe and incubated for 2 h at 40 °C in a humid chamber. The ileal tissues were washed twice in 1x wash buffer for 2 min at RT with agitation. Amplification and detection steps were performed using the RNAScope detection kit reagents. For the signal amplification, tissues were incubated in 50 μl of amplifier 1 solution (AMP 1) for 30 min at 40 °C and washed twice in 1x wash buffer for 2 min at RT with occasional agitation. Subsequently, the tissue was incubated in 50 μl of AMP 2 solution for 15 min at 40°C and then washed twice in 1x wash buffer for 2 min at RT with occasional agitation. After AMP 2 amplification, 50 μl of AMP 3 solution was added to the tissue preparations and incubated for 30 min at 40°C. The tissue was washed twice in 1x wash buffer for 2 min at RT with occasional agitation. Lastly, the tissue was incubated in 50 μl of AMP 4 solution and incubated for 15 min at 40°C prior to washing the tissue twice in 1x wash buffer for 2 min at RT with occasional agitation. The same protocol was followed for control experiments and dapB probe was used as the negative control probe (Supplementary Figure S1).

### Image acquisition and analysis

Multi-channel image acquisition was performed with a laser scanning confocal microscope 800 (Carl Zeiss Microscopy, North Ryde, NSW, Australia) using a 40x oil immersion objective lens. The primary and secondary antibodies used in this study are listed in Table 1.1 and Table 1.2 respectively. The samples were excited with Diode lasers at 561, 488 and 647 wavelengths. The pinhole diameter, detector gain and laser powers were optimised to obtain the highest pixel intensity at the same time eliminating pixel saturation. Consecutive Z stacks on the horizontal plane with a frame size of 1024x1024 pixels and a file-depth of 16 bits were captured at 1 µm intervals. Z-stacks together with tile scans were performed where necessary. The quantification was performed using Imaris 9.0 image analysis software (Bitplane, UK). Imaris software includes 3D reconstruction tools and statistical annotation for quantification purposes. Three-dimensional cell rendering was undertaken using high-resolution Z-stack fluorescent images generated using confocal microscopy. To create a detailed rendered surface, the “Create surface” tool in the Imaris software package was used. The minimum diameter of the surface was determined using the “Slice view” option. The interactive software histogram within the “Create surface” window was used to set a threshold (the minimum diameter of the neuronal surface) to exclusively include neuronal surfaces and to exclude background noise. Aggregated neurons within images were separated using the ‘Split surfaces” tool in Imaris. The Imaris software is additionally equipped with “Filter options” to label specific populations of surfaces either manually or automatically and these were utilized in the current study to identify specific cell populations in any given image. Since the RNAScope signal representing *Nlgn3* mRNA appears as puncta, the “spot” analysis tool in Imaris was used to create the 3D structure of individual mRNA molecules. The average diameter of spots detected was determined using “slice view” in Imaris. Once spots were generated, they were assigned as being located within the neurons using the “Split spots on to surface” tool. The Imaris 9.0 software additionally generates statistical data yielding the number of spots per surface. This option was used to identify the number of mRNA punctate signals within the neurons for the images analysed in the current study.

**Table 1.1.** Primary antibodies used for immunocytochemistry.

| **Primary antisera** | **Raised in** | **Dilution** | **Source** |
| --- | --- | --- | --- |
| Hu | Human | 1:5000 | Gift from Dr. V. Lennon |
| VIP | Rabbit | 1:1000 | Merck Millipore |
| ChAT | Goat | 1:100 | Chemicon |
| NOS | Sheep | 1:1000 | Gift from Dr P. Emson |
| Calretinin | Goat | 1:1000 | SWANT |

**Table 1.2.**
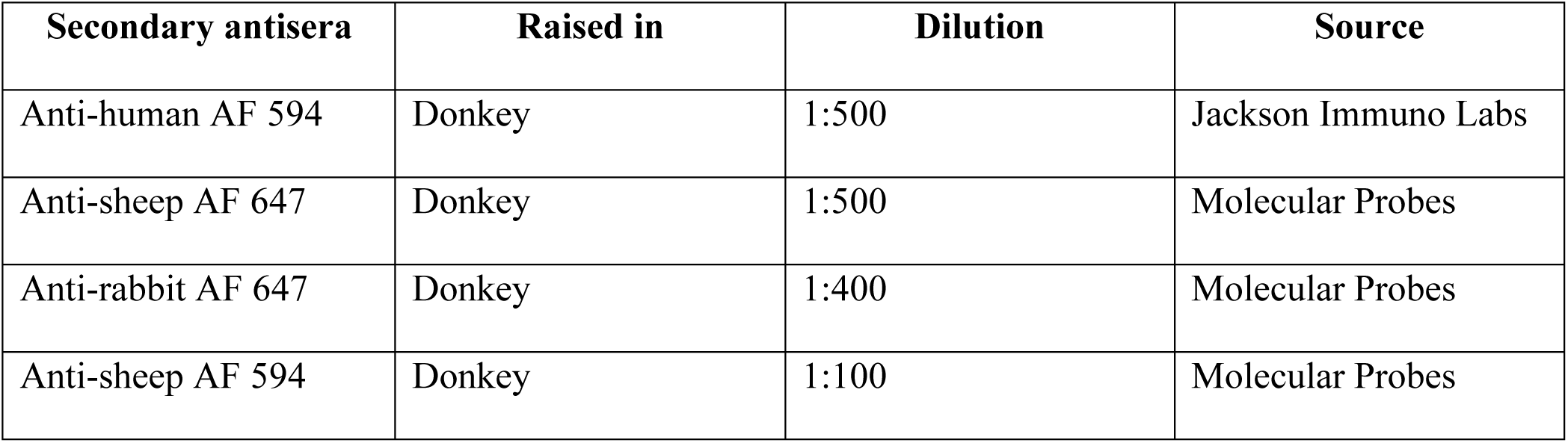
Secondary antibodies used for immunocytochemistry.

| Secondary antisera | Raised in | Dilution | Source |
| --- | --- | --- | --- |
| Anti-human AF 594 | Donkey | 1:500 | Jackson Immuno Labs |
| Anti-sheep AF 647 | Donkey | 1:500 | Molecular Probes |
| Anti-rabbit AF 647 | Donkey | 1:400 | Molecular Probes |
| Anti-sheep AF 594 | Donkey | 1:100 | Molecular Probes |

### Statistical analysis

Data were analysed using GraphPad Prism 8.4.1 (GraphPad software, USA). Frequency distribution analysis was performed to analyse the distribution of *Nlgn3* mRNA expression in enteric neurons and glia and a Kolmogorov-Smirnov test was conducted to determine statistical significance. To estimate the extent of overlap between two distributions for *Nlgn3* mRNA expression between genotypes, the magnitude of Kolmogorov-Smirnov D (D) was used.

## RESULTS

In this study we first determined the overall *Nlgn3* mRNA expression profile in each neuronal population using frequency distribution analysis. We then reported average *Nlgn3* mRNA copy numbers per cell in each of these cell populations.

### *Nlgn3* mRNA is expressed in the majority of distal ileal submucosal neurons

We first determined the expression of *Nlgn3* mRNA expression in all ileal submucosal neurons and cholinergic and non-cholinergic subpopulations using a dual-RNAScope/immunofluorescence and Imaris-based quantitative image analysis method. All submucosal neurons in distal ileal preparations were labelled with the pan-neuronal marker Hu (**Figure 1 A1-A6**). To analyse the distribution of *Nlgn3* mRNA expression in individual submucosal neurons, we assessed *Nlgn3* mRNA expression in a total of 1,985 submucosal neurons in 14 WT mice. Frequency distribution analysis of expression data acquired from within the cell somata revealed that 1724 of 1985 submucosal neurons (87%) expressed *Nlgn3* mRNA **(Figure 1 A7)**. In addition to *Nlgn3* mRNA in cell soma, our study also show the *Nlgn3 mRNA* expression in neuronal and glial fibers (Supplementary Figure S2).

**Figure 1.**
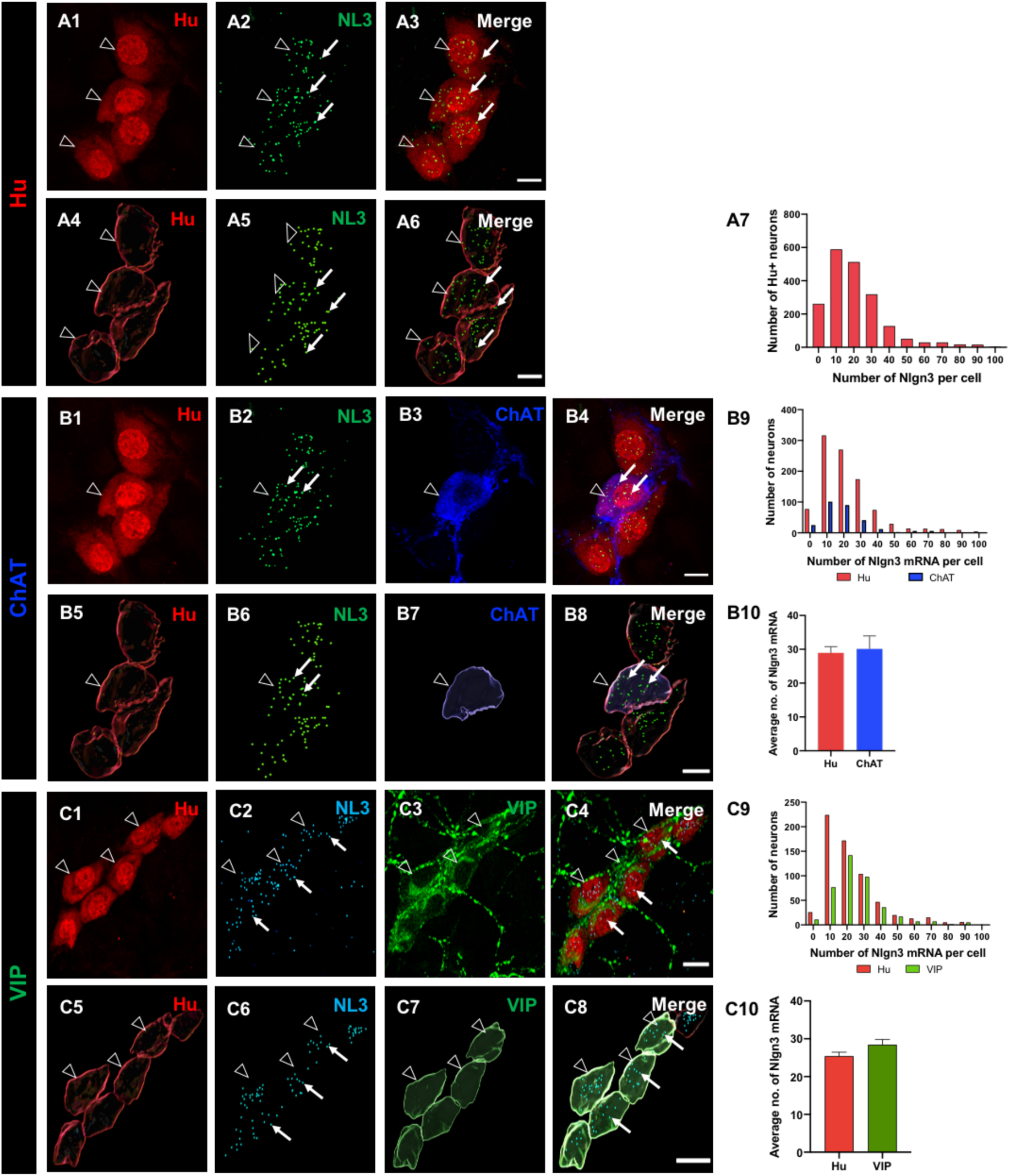
The distribution of *Nlgn3* mRNA in ileal submucosal neurons. Confocal images of (**A1**) submucosal neurons labelled with Hu pan neuronal marker (**A2**) *Nlgn3* mRNA expression (**A3**) *Nlgn3* mRNA expression in a submucosal ganglion. 3D reconstruction of (**A4**) Submucosal neurons (**A5**) *Nlgn3* mRNA (**A6**) *Nlgn3* mRNA-expressing submucosal neurons. (**A7**) Frequency distribution of *Nlgn3* mRNA expression in submucosal neurons. Confocal images of (**B1**) Submucosal ganglion (**B2**) *Nlgn3* mRNA expression (**B3**) ChAT neuron (**B4**) *Nlgn3* mRNA expression in an individual ChAT neuron. 3D structure of (**B5**) submucosal neurons (**B6**) *Nlgn3* mRNA (**B7**) a cholinergic submucosal neuron (**B8**) *Nlgn3* mRNA expression in an individual ChAT neuron. **B9**. The distribution of *Nlgn3* mRNA expression in cholinergic neurons is similar to that in total neurons in the submucosal plexus. **B10**. Average number of *Nlgn3* mRNA copies in cell soma in cholinergic neurons is similar to that in total neurons in the submucosal plexus. Confocal micrographs of (**C1**) submucosal neurons (**C2**) *Nlgn3* mRNA expression (**C3**) VIP expression in submucosal neurons (**C4**) *Nlgn3* mRNA expression in VIP neurons. 3D reconstruction of (**C5**) submucosal neurons (**C6**) *Nlgn3* mRNA expression (**C7**) VIP-expressing neurons (**C8**) *Nlgn3* mRNA expression in VIP-positive neurons. **C9**. Frequency distribution of *Nlgn3* mRNA expression in VIP submucosal neurons is significantly different to total submucosal neurons. **C10**. On average, VIP submucosal neurons express similar numbers of *Nlgn3* mRNA copies as observed in the total number of neurons in the submucosal plexus. Hu staining is indicated by open arrowheads and *Nlgn3* mRNA is indicated by arrows. Scale bar=10 µm

To assess for differential distribution patterns of *Nlgn3* mRNA within the submucosal plexus, *Nlgn3* mRNA expression levels in different submucosal neuron subpopulations were evaluated. The expression of *Nlgn3* mRNA in cholinergic neurons (labelled using ChAT) in 298 neurons from 7 WT mice was determined. Utilizing the frequency distribution analysis approach, we identified that the vast majority of cholinergic neurons express *Nlgn3* mRNA (274 of 298 neurons; 92%; **Figure 1 B1-B8**). The distribution profile of *Nlgn3* mRNA expression (copy number) in cholinergic neurons was similar to the *Nlgn3* mRNA distribution in Hu-positive submucosal neurons (D=0.08; p=0.09; **Figure 1 B9**). We also compared the expression in terms of *Nlgn3* mRNA copy numbers in cholinergic neurons. On average, *Nlgn3* mRNA-expressing cholinergic submucosal neurons contain 30.1 ± 3.8 copies of *Nlgn3* mRNA and Hu-positive submucosal neurons contain 29.0 ± 1.8 copies (n=298; n=1,014; ChAT and Hu positive neurons respectively; p=0.7; **Figure 1 B10)**.

Next, we measured the expression of *Nlgn3* mRNA in non-cholinergic (VIP-containing) submucosal neurons (**Figure 1 C1-C8**). A total of 411 VIP submucosal neurons from 7 WT mice were analysed. As shown by the frequency distribution analysis, the majority (374 of 411, 91%) of VIP-positive neurons express *Nlgn3* mRNA. The distribution profile of *Nlgn3* mRNA in VIP neurons is significantly different compared to the distribution of *Nlgn3* mRNA in Hu-positive submucosal neurons (D=0.16; p<0.0001) (**Figure 1 C9**). Despite the difference in distribution profile of *Nlgn3* mRNA, neurons that co-express *Nlgn3* mRNA and VIP contain an average of 28.5 ± 1.1 *Nlgn3* mRNA copies per neuron (n=411 neurons), similar to the number of *Nlgn3* mRNA copies expressed in total submucosal neurons overall (25.4 ± 0.7 copies per neuron, n=644 neurons; p=0.06) **(Figure 1 C10)**.

### Myenteric neurons express *Nlgn3* mRNA

The subcellular distribution of *Nlgn3* mRNA in the distal ileum was analysed in a total of 3,788 myenteric neurons from 14 WT mice. The majority (3295 of 3788, 87%) of myenteric neurons expressed *Nlgn3* mRNA within the cell soma. The remaining 13% (492 neurons) of myenteric neurons did not express detectable levels of *Nlgn3* mRNA (**Figure 2 A1-A6**). Overall, myenteric neurons expressing *Nlgn3* mRNA contained an average of 16.8 ± 0.2 copies of *Nlgn3* mRNA in the cell soma **(Figure 2 A7)**.

**Figure 2.**
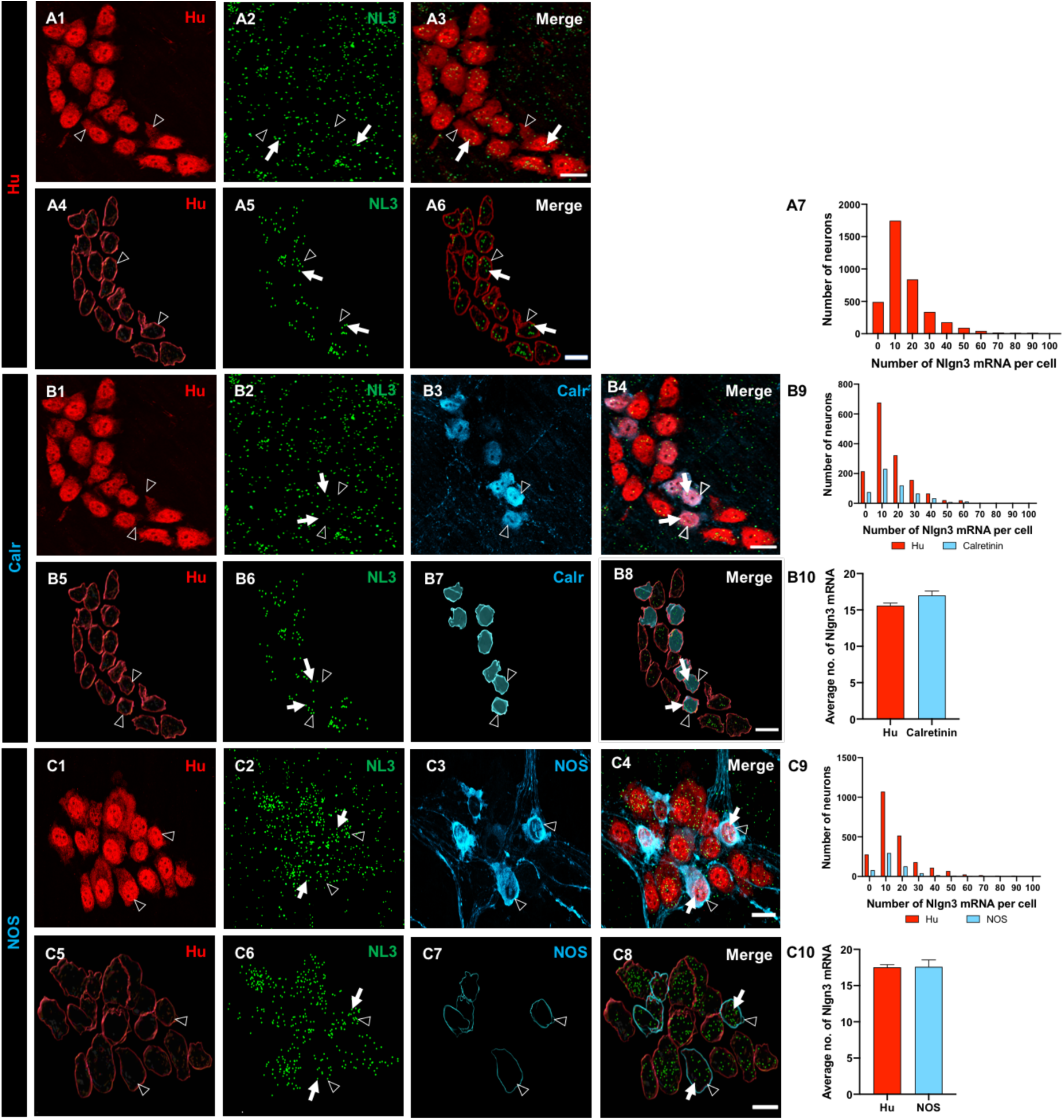
Expression and the distribution of *Nlgn3* mRNA in distal ileal myenteric neurons. **A1**. Myenteric neurons labelled using Hu by immunofluorescence. **A2**. *Nlgn3* mRNA expression was detected using RNAScope. **A3**. *Nlgn3* mRNA expression in myenteric neurons. The 3D structure of (**A4**) myenteric neurons (**A5**) *Nlgn3* mRNA (**A6**) *Nlgn3* mRNA containing myenteric neurons. **A7**. Frequency distribution of *Nlgn3* expression in myenteric neurons. Confocal micrographs of (**B1**) myenteric neurons labelled with the pan-neuronal marker, Hu (**B2**) *Nlgn3* mRNA (**B3)** Calretinin positive myenteric neurons. **B4**. Expression of *Nlgn3* mRNA in calretinin neurons. The 3D rendering of (**B5**) myenteric neurons (**B6**) *Nlgn3* mRNA distribution in the myenteric plexus, (**B7**) calretinin expressing neurons and (**B8**) *Nlgn3* mRNA expression in calretinin-expressing myenteric neurons. **B9**. The frequency distribution of *Nlgn3* mRNA expression in calretinin-positive myenteric neurons is similar to total myenteric neurons. **B10**. Calretinin-expressing neurons contain similar copy numbers of *Nlgn3* mRNA to that of myenteric neurons overall. Triple labelling of (**C1**) myenteric neurons, (**C2**) *Nlgn3* mRNA and (**C3**) NOS expressing myenteric neurons. **C4**. The expression of *Nlgn3* mRNA in NOS-containing myenteric neurons. The 3D structure of (**C5**) myenteric neurons, (**C6**) *Nlgn3* mRNA, (**C7**) NOS-expressing myenteric neurons and (**C8**) *Nlgn3* mRNA in NOS-containing neurons. **C9**. The frequency distribution of *Nlgn3* mRNA in NOS-containing neurons is similar to that in the total population (i.e. Hu-positive neurons). **C10**. The average number of *Nlgn3* mRNA copies in myenteric NOS-containing neurons is similar to the total number of myenteric neurons. Hu staining is indicated by open arrowheads and *Nlgn3* mRNA labelling is indicated by arrows. Scale bar=10 µm.

To understand whether *Nlgn3* mRNA is differentially expressed between neuronal subtypes within the myenteric plexus, we labelled subpopulations of myenteric neurons using immunofluorescence for cell type-specific markers. Expression data showed that 473 of 550 (86%) of calretinin-immunoreactive neurons express *Nlgn3* mRNA. Frequency distribution analysis of *Nlgn3* mRNA expression in calretinin neurons showed a similar profile to *Nlgn3* mRNA expression in the total myenteric neuronal population (Hu-positive neurons; n=1484, calretinin-positive neurons; n=550; D=0.04; p=0.2). There was no significant difference in *Nlgn3* mRNA copy numbers in calretinin neurons compared to Hu-positive myenteric neuron populations (calretinin positive neurons; 17.0 ± 0.6 mRNA copies, Hu-positive neurons;15.6 ± 0.3 mRNA copies; p=0.2; **Figure 2 B1-B10)**.

Co-labelling of NOS-expressing myenteric neurons with RNAScope revealed that 515 of 593 (87%) of NOS neurons express *Nlgn3* mRNA in the cell soma. The frequency distribution of *Nlgn3* mRNA in NOS-containing neurons was similar to that of Hu-positive neurons in the myenteric plexus (NOS positive neurons; n=593, Hu positive neurons; n=2304, p=0.09). An individual NOS-containing neuron contained an average of 17.6 ± 0.9 *Nlgn3* mRNA copies, n=593 neurons), which was similar to the number of copies expressed in Hu-positive myenteric neurons (17.5 ± 0.4 *Nlgn3* mRNA copies, n=2304 neurons; D=0.05; p=0.09; **Figure 2 C1-C10**).

### Enteric glia express *Nlgn3*

Recent studies revealed that *Nlgn3* is also expressed in non-neuronal (enteroendocrine) cells in the intestine (Bohorquez et al., 2015) but whether *Nlgn3* is expressed in enteric glia has not been assessed. We therefore labeled submucosal glia using an antiserum against the calcium-binding protein, S100β and assessed for the co-presence of *Nlgn3* mRNA (**Figure 3 A1-A4**). Overall, we found that 64% (144 of a total of 226 cells in 6 individual WT mice) of submucosal glial cells in the mouse ileum express *Nlgn3* mRNA (**Figure 3 A1-A9**). Similarly, *Nlgn3* mRNA expression in ileal myenteric glial cells (1203 glial cells in 4 WT mice) was assessed (**Figure 3 B1-B8**). In contrast with the submucosal plexus, only 37% (445 of 1203 cells) of myenteric glial cells expressed *Nlgn3* mRNA in the mouse ileum (**Figure 3 B9**).

**Figure 3.**
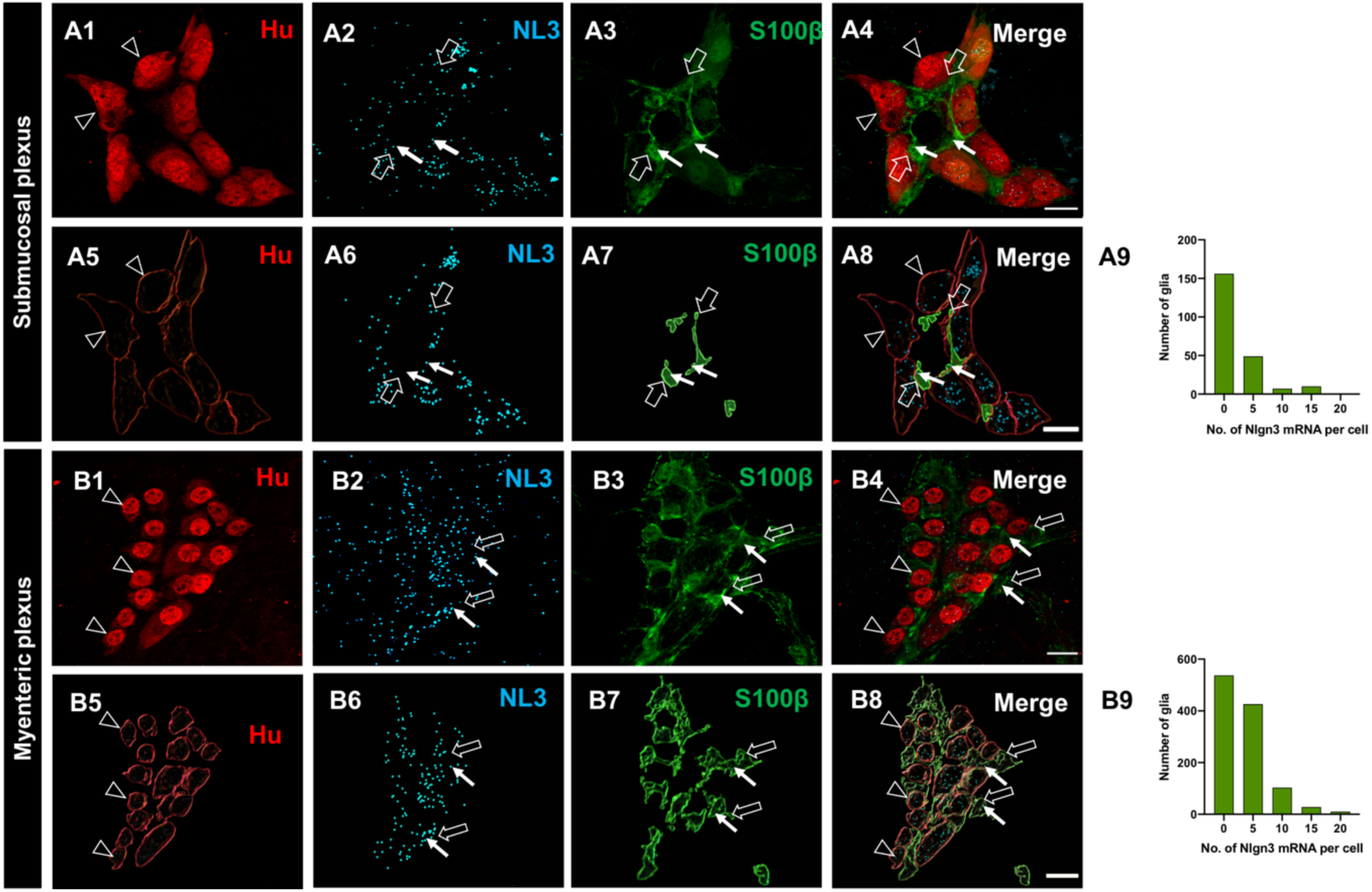
The expression and distribution of *Nlgn3* mRNA in enteric glia. Confocal micrographs of (**A1**) submucosal neurons labelled with the pan-neuronal marker Hu (**A2**) *Nlgn3* mRNAexpression in the submucosal plexus (**A3**) submucosal glia labelled with S100β (**A4**) *Nlgn3* mRNA in submucosal glia. Imaris-based 3D rendering of (**A5**) submucosal neurons (**A6**) *Nlgn3* mRNA (**A7**) submucosal glia (**A8**) *Nlgn3* mRNA expression in submucosal glia. **A9**. The frequency distribution analysis demonstrates that *Nlgn3* mRNA is expressed in the majority of submucosal glial cells. Triple labelling of (**B1**) myenteric neurons using Hu pan-neuronal marker (**B2**) *Nlgn3* mRNA (**B3**) myenteric glia using S100β. **B4**. The expression of *Nlgn3* mRNA in myenteric glia. 3D rendering of (**B5**) myenteric neurons (**B6**) *Nlgn3* mRNA (**B7**) myenteric glia (**B8**) *Nlgn3* mRNA expression in myenteric glia. **B9**. Frequency distribution analysis displayed that the majority of myenteric glia do not express *Nlgn3* mRNA. Hu staining is indicated by the open arrowhead, *Nlgn3* mRNA is indicated by the filled arrow and submucosal glia are labelled with open arrows. Scale bar=10 µm

### The R451C mutation reduces *Nlgn3* mRNA expression in cholinergic submucosal neurons but not in VIPergic neurons

We next investigated *Nlgn3* mRNA expression in wild type and *Nlgn3*^R451C^ mutant mouse ileum using dual RNAScope and immunocytochemistry plus Imaris-based quantification (**Figure 4 A1-A6**). Frequency distribution analysis of *Nlgn3* mRNA expression in the submucosal plexus showed that *Nlgn3* mRNA expression is significantly different in *Nlgn3*^R451C^ mutant mice compared to WT (1985 neurons in 14 WT and 1300 neurons in 10 *Nlgn3*^R451C^ mice were analysed; D=0.15; p<0.0001; **Figure 4 A7**). In *Nlgn3*^R451C^ mutant mice, submucosal neuronal cell bodies express 21.6 ± 0.8 (n=1300 neurons) copies of *Nlgn3* mRNA, indicating that significantly less *Nlgn3* mRNA is present compared with in WT mice (24.8 ± 1.0 copies, n=1985 neurons), p<0.000; **Figure 4 A8**).

**Figure 4.**
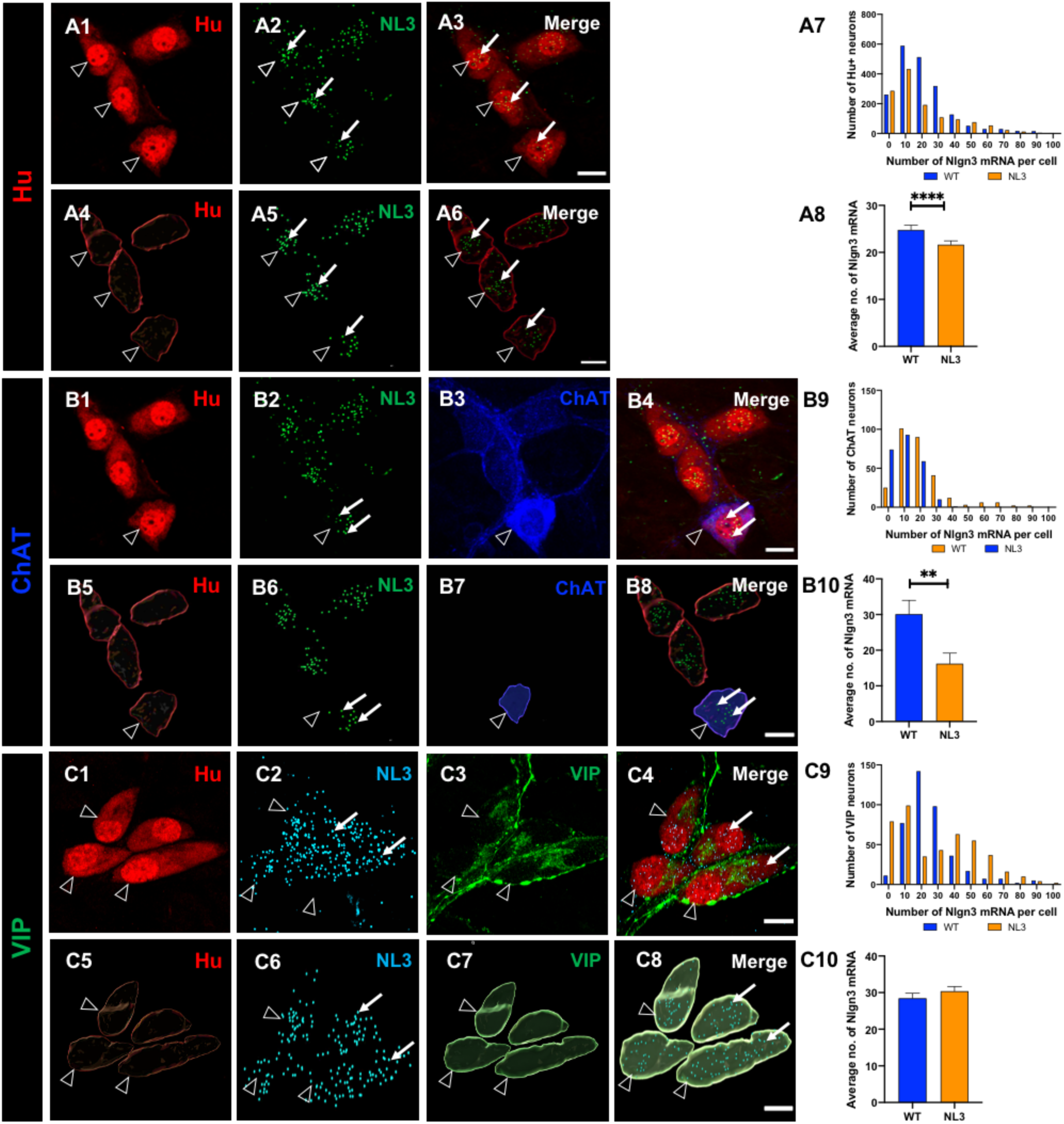
Expression of *Nlgn3* mRNA in the ileal submucosal plexus of *Nlgn3*^R451C^ mutant mice. Confocal micrographs of (**A1**) Submucosal neurons (**A2**) *Nlgn3* mRNA expression (**A3**) *Nlgn3* mRNA expression in submucosal neurons. The 3D structure of (**A4**) submucosal neurons (**A5**) *Nlgn3* mRNA (**A6**) *Nlgn3* mRNA expression in submucosal neurons. **A7.** The distribution of *Nlgn3* mRNA in *Nlgn3*^R451C^ mutant mice is significantly different compared to WT. **A8**. *Nlgn3*^R451C^ mutant mice express less *Nlgn3* mRNA in submucosal neurons compared to WT. Triple labelling of (**B1**) Submucosal neurons with Hu pan-neuronal marker (**B2**) *Nlgn3* mRNA (**B3**) cholinergic neurons with ChAT antibody. **B4**. The expression of *Nlgn3* mRNA in cholinergic neurons. 3D rendering of (**B5**) Hu neurons (**B6**) *Nlgn3* mRNA (**B7**) ChAT neuron (**B8**) *Nlgn3* mRNA expression in ChaT neurons. **B9**. The frequency distribution of *Nlgn3* mRNA in cholinergic submucosal neurons in *Nlgn3*^R451C^ mutant mice compared to WT **B10**. Cholinergic neurons express significantly less *Nlgn3* mRNA in *Nlgn3*^R451C^ mutants compared to WT. Confocal micrographs of (**C1**) Submucosal neurons labelled with Hu pan-neuronal marker (**C2**) *Nlgn3* mRNA labelled with RNAScope (**C3**) Non-cholinergic neurons labelled with VIP (**C4**) *Nlgn3* expression in non-cholinergic submucosal neurons. 3D reconstruction of (**C5**) submucosal neurons (**C6**) *Nlgn3* mRNA (**C7**) VIP neurons (**C8**) *Nlgn3* mRNA in VIP neurons. **C9**. The frequency distribution of *Nlgn3* mRNA in non-cholinergic submucosal neurons in *Nlgn3*^R451C^ mutant mice compared to WT. **C10**. *Nlgn3*^R451C^ mice express similar numbers of *Nlgn3* mRNA copies in non-cholinergic neurons compared to WT in the submucosal plexus. *Nlgn3* mRNA is indicated by the filled arrow and submucosal glia are labelled with open arrows **p<0.01, ****p<0.0001, Scale bar=10 µm

We then characterised the expression of *Nlgn3* mRNA in different submucosal neuronal populations (**Figure 4 B1-B8**). Of a total of 241 ChAT neurons in 7 *Nlgn3*^R451C^ mice, only 69% (166 neurons) of neurons expressed *Nlgn3* mRNA. The frequency distribution analysis revealed that the distribution of *Nlgn3* mRNA in ChAT neurons in mutant mice is significantly different from WT (n=298 neurons in 5 WT mice, n=241 neurons in *Nlgn3*^R451C^ mice; D=0.32; p<0.0001; **Figure 4 B9**). Specifically, in *Nlgn3*^R451C^ mutant mice there is a significant reduction in *Nlgn3* mRNA expression in ChAT neurons compared to WT (WT; 30.1 ± 3.8 copies, n=298, *Nlgn3*^R451C^ mutant mice; 16.2 ± 2.9 copies, n=241, p=0.006; **Figure 4 B10**).

The impact of the R451C mutation on *Nlgn3* mRNA production in VIP-expressing neurons was also evaluated (**Figure 4 C1-C8**). The frequency distribution of the *Nlgn3* mRNA copy number distribution in VIP neurons in ileal tissue from *Nlgn3*^R451C^ mice (n=623 neurons in 7 WT mice were analysed as well as n=449 neurons in 5 *Nlgn3*^R451C^ mice), differs significantly from WT (D=0.15; p<0.0001; **Figure 4 C9**). In WT mice, 91% (566 of 623) of VIP-expressing neurons express *Nlgn3* mRNA and in mutant mice, only 82% (409 of 449) of VIP-containing submucosal neurons express *Nlgn3* mRNA in the cell soma. In VIP-expressing neurons, *Nlgn3* mRNA expression levels showed a trend to increase in mutant mice compared to WT (27.1 ± 1.1; 30.4 ± 1.2 *Nlgn3* mRNA copies per cell; n=623 and 449 cells, WT and mutant mice respectively; p=0.05; **Figure 4 C10**).

### The R451C mutation decreases *Nlgn3* mRNA expression in myenteric neurons

We evaluated the impact of the R451C mutation on *Nlgn3* mRNA expression distribution profiles and mRNA copy number in the ileal myenteric plexus (**Figure 5 A1-A6**). Using the same approach, we compared *Nlgn3* mRNA expression in Hu, Calretinin and NOS-immunoreactive myenteric neurons in WT and *Nlgn3*^R451C^ mutant mice. We found that the frequency distribution of *Nlgn3* mRNA expression in the myenteric plexus was significantly different between WT and *Nlgn3*^R451C^ mice (D=0.16; p<0.0001; **Figure 5 A7**). As expected, *Nlgn3* mRNA copy numbers in myenteric neurons were significantly reduced in *Nlgn3*^R451C^ mutant mice compared to WT (WT; 16.8 ± 0.3 mRNA copies per cell, *Nlgn3*^R451C^ mice; 11.3 ± 0.2 mRNA copies per cell, n=3,788, n=2,825, WT and *Nlgn3*^R451C^ mutant mice respectively; p<0.0001; **Figure 5 A8**).

**Figure 5.**
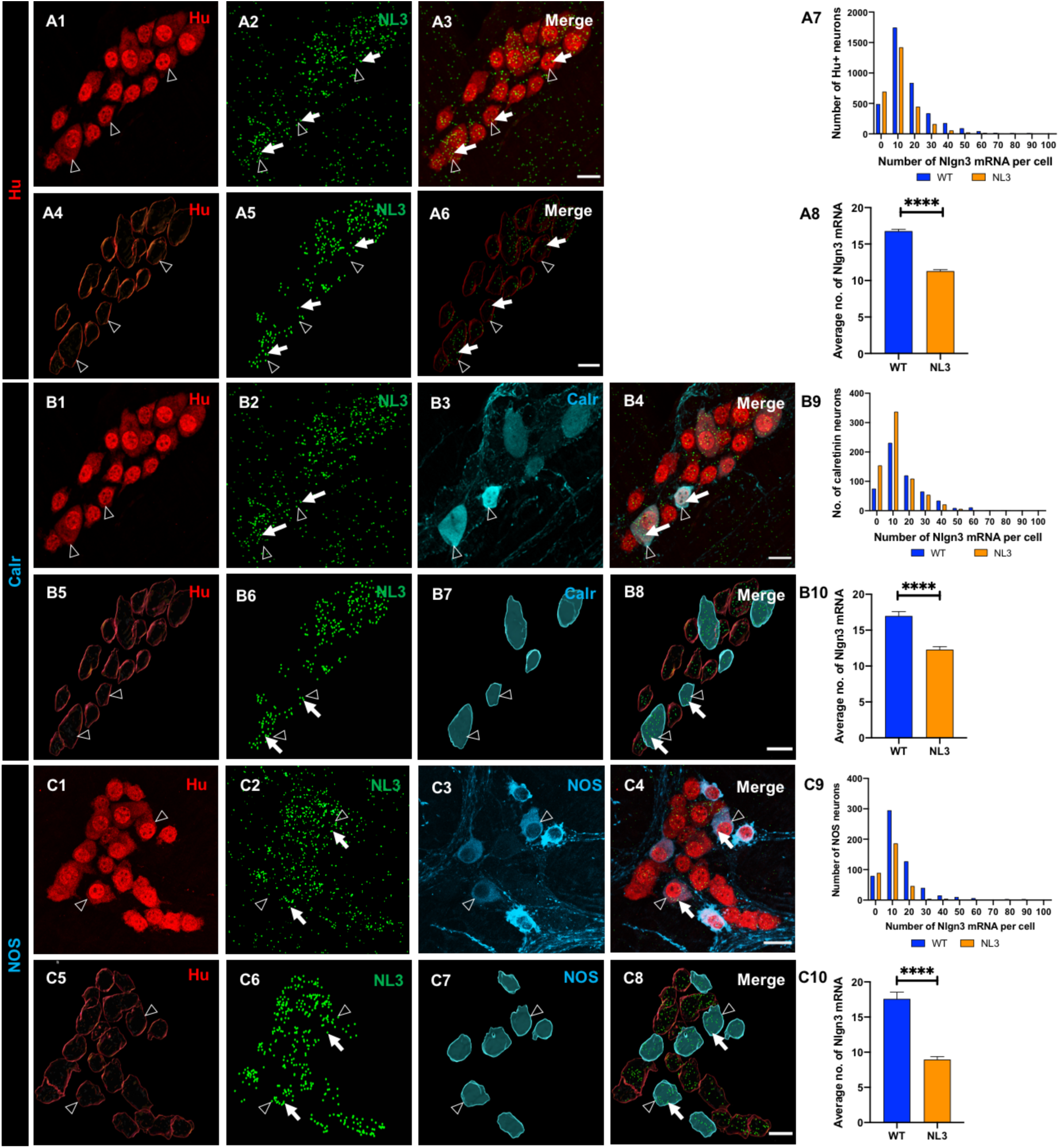
Effects of the *Nlgn3* R451C mutation on *Nlgn3* mRNA expression in myenteric neurons. Confocal images of (**A1**) myenteric neurons (**A2**) *Nlgn3* mRNA expression (**A3**) *Nlgn3* mRNA expression in myenteric neurons. The 3D structure of (**A4**) myenteric neurons (**A5)** *Nlgn3* mRNA (**A6)** *Nlgn3* mRNA expression in myenteric neurons. **A7**. The frequency distribution of *Nlgn3* mRNA in *Nlgn3*^R451C^ mutant mice compared WT **A8**. In the *Nlgn3*^R451C^ mouse ileum, myenteric neurons express fewer copies of *Nlgn3* mRNA compared to WT. Triple labelling of (**B1**) myenteric neurons using Hu pan-neuronal marker (**B2**) *Nlgn3* mRNA (**B3**) calretinin neurons. **B4**. The expression of *Nlgn3* mRNA in calretinin expressing neurons in *Nlgn3*^R451C^ mutant mice. 3D reconstruction of (**B5**) myenteric neurons (**B6**) *Nlgn3* mRNA (**B7**) calretinin expressing neurons and (**B8**) *Nlgn3* mRNA expression calretinin neurons in *Nlgn3*^R451C^ mutant mice. **B9**. The frequency distribution of *Nlgn3* mRNA in calretinin positive myenteric neurons in *Nlgn3*^R451C^ mutant mice and WT. **B10**. The *Nlgn3* R451C mutation reduces the *Nlgn3* mRNA expression in mutant mice compared to WT. Triple labelling of (**C1**) myenteric neurons (**C2**) *Nlgn3* mRNA (**C3**) NOS neurons (**C4**) *Nlgn3* mRNA in NOS expressing myenteric neurons. 3D rendering of (**C5**) myenteric neurons (**C6**) NOS positive neurons (**C7**) *Nlgn3* mRNA and (**C8**) *Nlgn3* mRNA expressing NOS neurons. **C9**. The frequency distribution of *Nlgn3* mRNA expression in NOS containing neurons in *Nlgn3*^R451C^ mutant mice is significantly different compared to WT **C10**. In *Nlgn3*^R451C^ mutant mice, NOS neurons contain less *Nlgn3* mRNA compared to WT. *Nlgn3* mRNA is indicated by the filled arrow and submucosal glia are labelled with open arrows, ****p<0.0001, Scale bar=10 µm.

We also assessed for differences in *Nlgn3* mRNA expression in calretinin-immunoreactive myenteric neurons (**Figure 5 B1-B8**). The frequency distribution of *Nlgn3* mRNA in calretinin neurons in *Nlgn3*^R451C^ mice is significantly different from the WT (D=0.16, p<0.0001, **Figure 5 B9**). In WT, 86% (473 of 550) of calretinin neurons express *Nlgn3* mRNA, of the total calretinin positive cellular population in *Nlgn3*^R451C^ mutant mice, 78% of neurons co-express *Nlgn3* mRNA. Significantly fewer *Nlgn3* mRNA copies were detected in calretinin-immunoreactive neurons in *Nlgn3*^R451C^ mutant mice compared to WT (WT;17.0 ± 0.6 mRNA copies, n=550 neurons*, Nlgn3*^R451C^ mice; 12.3 ± 0.42 mRNA copies, n=685 neurons; 0.16 p<0.0001; **Figure 5 B10**).

It has been reported that *Nlgn3*^R451C^ mice have an increased proportion of NOS neurons in the myenteric plexus of the jejunum and caecum (Hosie et al., 2019, Sharna et al., 2020), suggesting that NLGN3 might play an important role in NOS signalling in the myenteric plexus. Therefore, we analysed the impact of the R451C mutation on *Nlgn3* mRNA expression in NOS-immunoreactive myenteric neurons (**Figure 5 C1-C8**). The frequency distribution of *Nlgn3* mRNA distribution in NOS immunoreactive neurons in *Nlgn3*^R451C^ mice differs significantly from WT (WT; n=593 neurons, *Nlgn3*^R451C^ mice; n=331, D=0.26; p<0.0001, **Figure 5 C9**). In WT, about 87% of NOS-expressing neurons (i.e., 515 of 593 neurons) express *Nlgn3* mRNA. Of a total of 331 NOS-immunoreactive neurons assessed in *Nlgn3*^R451C^ mutant mice, about 73% (241 neurons) co-express *Nlgn3* mRNA. *Nlgn3* mRNA expression levels in NOS-immunoreactive neurons in the myenteric plexus were significantly reduced in mutant mice compared to WT (WT; 17.6 ± 0.9 mRNA copies per cell, n=593 neurons, *Nlgn3*^R451C^ mutant mice; 9.0 ± 0.4 mRNA copies per cell, n=331 neurons, p<0.0001; **Figure 5 C10**).

### *Nlgn3* mRNA is reduced in ileal myenteric glia in *Nlgn3*^R451C^ mice

As described earlier, *Nlgn3* mRNA is expressed in both submucosal and myenteric glia in the WT mouse ileum. To determine if the R451C mutation alters glial expression patterns of *Nlgn3* mRNA, expression levels of *Nlgn3* mRNA were assessed in 226 and 173 glial cells of the ileal submucosal plexus in WT (n=5) and *Nlgn3*^R451C^ (n=5) mutant mice, respectively (**Figure 6 A1-A8**). The frequency distribution data of *Nlgn3* mRNA expression in glial cells show similar levels of *Nlgn3* mRNA expression in WT and *Nlgn3*^R451C^ submucosal glia (n=226 neurons in 6 WT mice, n=173 neurons in *Nlgn3*^R451C^ mice; D=0.12; p=0.09, **Figure 6 A9**). In submucosal glia, there was a trend for an increase in mRNA copy number in *Nlgn3*^R451C^ mutants (2.8 ± 0.3 mRNA copies in n=226 glial cells; 5.0 ± 1.2 mRNA copies in n=173 glial cells; WT and *Nlgn3*^R451C^ mice respectively, p=0.05; **Figure 6 A10**).

**Figure 6.**
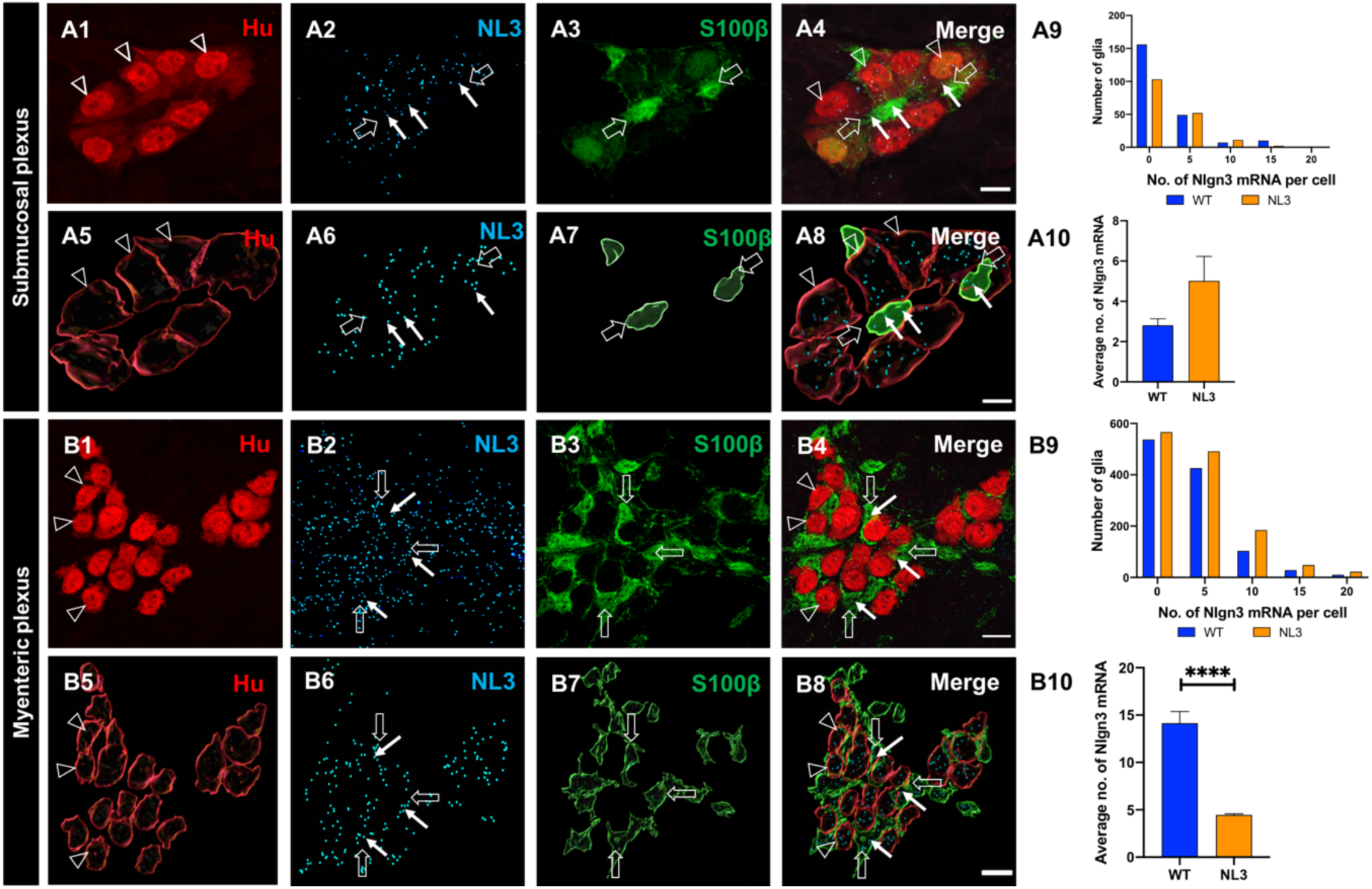
Effects of *Nlgn3* R451C mutation on *Nlgn3* mRNA expression in enteric glia. Confocal images of (**A1**) submucosal neurons label with Hu (**A2**) *Nlgn3* mRNA expression in the submucosal plexus (**A3**) submucosal glia labelled with S100β (**A4**) *Nlgn3* mRNA expression in submucosal glia. The 3D structure of (**A5**) submucosal neurons (**A6**) *Nlgn3* mRNA (**A7**) submucosal glia (**A8**) *Nlgn3* mRNA expression in submucosal glia. **A9**. Frequency distribution of *Nlgn3* mRNA in submucosal glia in *Nlgn3*^R451C^ mutant mice compared to WT. **A10**. The copy number expression of *Nlgn3* mRNA per cell in WT compared to *Nlgn3*^R451C^ mutant mice. Immunofluorescent labelling of (**B1**) myenteric neurons (**B2**) *Nlgn3* mRNA (**B3**) myenteric glia (**B4**) *Nlgn3* mRNA localization in myenteric glia in *Nlgn3*^R451C^ mutant mice. 3D reconstruction of (**B5**) myenteric neurons (**B6**) *Nlgn3* mRNA (**B7**) myenteric glia (**B8**) *Nlgn3* mRNA expression in myenteric glia in *Nlgn3*^R451C^ mice. **B9**. Frequency distribution of *Nlgn3* mRNA expression in *Nlgn3*^R451C^ mutant mice is significantly different to WT mice. **B10**. Mutant mice express significantly fewer *Nlgn3* mRNA copies in myenteric glia compared to WT. ****p<0.0001, Scale bar=10 µm

We also compared *Nlgn3* mRNA expression in myenteric glia in WT and *Nlgn3*^R451C^ mutant mice. In *Nlgn3*^R451C^ mutant mice, of a total of 1318 glial cells, only 758 (57%) myenteric glia express *Nlgn3* mRNA (**Figure 6 B1-B8**) and the frequency distribution of *Nlgn3* mRNA in myenteric glia in mutant mice is significantly different to WT (n=1203 in 5 WT mice, n=1318 in 5 *Nlgn3*^R451C^ mice; D=0.07; p=0.001, **Figure 6 B9**). *Nlgn3* mRNA expression is reduced in myenteric glia in *Nlgn3*^R451C^ mutant mice compared to WT (WT; 14.2 ± 1.2 mRNA copies/cell, n=1203, *Nlgn3*^R451C^ mutant mice; 4.4 ± 0.1 mRNA copies/cell, n=1,318, p<0.0001, **Figure 6 B10**).

## DISCUSSION

In this study, we identified that most submucosal and myenteric neurons express *Nlgn3* mRNA in the cell soma of the mouse ileum. For the first time, this study revealed that *Nlgn3* mRNA is expressed in ileal submucosal and myenteric glia. In addition, we report that the autism-associated *Nlgn3* R451C mutation reduces *Nlgn3* mRNA expression in both submucosal and myenteric neurons as well as myenteric glia in the mouse ileum.

### Most enteric neurons express *Nlgn3* mRNA

Here we show that *Nlgn3* mRNA is expressed within the cell soma in most enteric neurons. Although some studies suggest the importance of NLGN3 for proper GI function (Leembruggen et al., 2019, Hosie et al., 2019), the cellular localization of NLGN3 in the ENS is unclear. Therefore, we conducted this study to understand the cellular localization of *Nlgn3* mRNA in order to co-relate the NLGN3 mediated mechanistic and functional involvement in regulating GI functions.

We reveal that most cholinergic submucosal neurons also contain *Nlgn3* mRNA. Within the enteric neural circuity, acetylcholine is the primary excitatory neurotransmitter utilised by cholinergic secretomotor neurons, excitatory muscle motor neurons, ascending interneurons, descending interneurons and intrinsic sensory neurons (Brookes, 2001, Grider, 2003, Gwynne and Bornstein, 2007, Brookes et al., 1991). Although there is no available evidence of NLGN3 expression in the cholinergic system of the central or peripheral nervous systems, expression of other NLGN subtypes in cholinergic synapses has been reported. For example, NLGN2 is expressed at the postsynaptic membrane of cholinergic synapses in the mouse brain (Takacs et al., 2013) and NLGN1 is present in cholinergic neurons in the chick ciliary ganglion (Rosenberg et al., 2010). In addition, NLGN3 and other NLGN isoforms can be colocalized in the same synapse (Budreck and Scheiffele, 2007). Based on this evidence, NLGN3 might be co-expressed with NLGN1 and NLGN2 in cholinergic synapses in the submucosal plexus. Since virtually all submucosal neurons receive synaptic inputs via cholinergic synapses (Foong et al., 2014), *Nlgn3* is likely present in cholinergic synapses in the submucosal plexus.

We also revealed that *Nlgn3* mRNA is expressed in VIP-expressing submucosal neurons in the mouse ileum, suggesting that NLGN3 might play a role in VIP signalling in the ENS. In the submucosal plexus, VIP is a primary neurotransmitter of secretomotor neurons and stimulates intestinal secretion (Foong et al., 2014, Mongardi Fantaguzzi et al., 2009). In contrast, VIP is expressed in excitatory neurons in the myenteric plexus. VIP also acts as a co-transmitter alongside ACh and NO in a subset of descending interneurons in the mouse and guinea pig ileum (Qu et al., 2008, Sang and Young, 1996, Costa et al., 1996). These findings open a novel avenue since the expression profiles of NLGN3 or other NLGN isoforms in VIP-containing synapses have not been reported to date in either the CNS or the peripheral nervous system.

One of the major findings of this study is that most NO-containing enteric neurons express *Nlgn3* mRNA in the cell soma. Nitric oxide (NO) is the predominant inhibitory neurotransmitter to the smooth muscle in the ENS (Qu et al., 2008, Sang and Young, 1996) and also expressed in descending interneurons together with other neurotransmitters (Young et al., 1995). Although the expression pattern of *Nlgn3* in NOS-expressing neurons is unknown, studies conducted in the *Nlgn3*^R451C^ mutant mice revealed that this mutation increases the number of NOS neurons in the myenteric plexus (Hosie et al., 2019). Based on this evidence, *Nlgn3* may contribute to NOS signalling in the ENS and we suggest that NLGN3 protein is likely expressed in inhibitory and descending interneurons in the myenteric plexus and contributes to NOS signalling in the mouse ENS. *Nlgn3* mRNA is also expressed in calretinin-expressing myenteric neurons in the mouse ileum. In the ENS, calretinin is expressed in intrinsic sensory neurons, interneurons and excitatory motor neurons innervating the smooth muscle layer (Sang and Young, 1996, Qu et al., 2008). Collectively, these data suggest that *Nlgn3* is expressed in almost all enteric neuronal types and therefore may play a functional role in synaptic transmission between different classes of enteric neurons (including secretory motor neurons, intrinsic sensory neurons, interneurons and muscle motor neurons).

### The R451C mutation reduces *Nlgn3* mRNA expression in the ENS

Mice expressing the *Nlgn3* R451C mutation display faster small intestinal transit and increased numbers of myenteric neurons alongside GABA_A_ receptor-mediated colon dysmotility (Hosie et al., 2019). But how this mutation affects NLGN3 expression in the enteric nervous system is not understood. Here we show that the R451C mutation decreases *Nlgn3* mRNA expression in both neurons and glia in the mouse ileal submucosal plexus. For example, cholinergic submucosal neurons in mutant mice contain reduced *Nlgn3* mRNA copy numbers compared to WT. These changes could potentially alter cholinergic signalling in the submucosal plexus which could affect cholinergic neuronal mediated functions such as secretion, absorption and mucosal barrier functions. Although there are no reports available on changes to the cholinergic system of the ENS in ASD, CNS studies showed that cholinergic neurons are associated with the pathophysiology of autism (Perry et al., 2001, Deutsch et al., 2010). Specifically, ASD patient basal forebrain tissues show altered cholinergic neuronal numbers, size and structure (Kemper and Bauman, 1998). In addition, a decreased plasma concentration of choline, a precursor for acetylcholine has been reported in ASD patients (Sokol et al., 2002, Wenk and Hauss-Wegrzyniak, 1999) and reduced levels of hippocampal cytosolic choline have been correlated with autism severity (Sokol et al., 2002). In the GI tract, NLGN3-mediated alterations to the cholinergic system could induce GI dysfunction in patients with neurological disorders such as ASD, however further studies are required to determine effects of the R451C mutation on cholinergic signalling in the ENS.

In the myenteric plexus of the distal ileum, the R451C mutation reduces *Nlgn3* mRNA expression levels in neurons. Specifically, *Nlgn3* mRNA expression is substantially reduced in both calretinin and NOS-immunoreactive neuronal populations. Although the contribution of calretinin neurons to ASD or ASD-related GI pathophysiology is unclear, the significance of calretinin signalling in other ENS-related diseases has been highlighted (Barshack et al., 2004). As mentioned, an increased proportion of NOS-expressing neurons has previously been observed in *Nlgn3^R451C^* mice (Hosie et al., 2019) therefore reduced levels of *Nlgn*3 expression might contribute to altered NOS neuronal signalling in the ENS in the *Nlgn3^R451C^* mouse model of autism.

### The R451C mutation dramatically reduces the expression of *Nlgn3* mRNA in enteric glia

For the first time, we reveal that *Nlgn3* mRNA is expressed in enteric glia. Specifically, we found that majority of enteric glia express *Nlgn3* mRNA therefore, we suggest that NLGN3 expressed in enteric glia might be involved in modulating glial-neuron synaptic activity in the ENS.

Enteric glia play a major role in ENS-mediated GI functions including mucosal secretion, intestinal permeability, mucosal sensation, GI motility and immune responses (Gulbransen and Sharkey, 2012, Grubisic et al., 2018) in concert with enteric neurons. Enteric glia synapse onto enteric neurons to regulate ENS-coordinated gut functions (Sharkey, KA., 2015. Enteric glial-neuronal associations revealed by immunocytochemistry highlight that enteric glia are in close contact with nerve fibers and varicose release sites (Boesmans et al., 2013). Evidence for neuro-glia communication also comes from live imaging experiments reporting the presence of several signalling pathways. A number of Ca^2+^ imaging studies indicate that neuron-glia communication is regulated by neurally released purines which activate purinergic receptors on enteric glia (Boesmans et al., 2013, Gomes et al., 2009, Gulbransen and Sharkey, 2009). In the submucosal plexus, neuron-glial transmission occurs via P2Y_1_ and P2Y_4_ receptors and via muscarinic receptors (ref). In addition, neuronal ATP release via the pannexin-1 channel also influences neuro-glia interactions (Wang and Dahl, 2018, Hanstein et al., 2016). Based on this evidence, we propose that NLGN3 expressed by both enteric neurons and glia modulate neuro-glia signalling pathways. This hypothesis is well-established in the mouse brain where RNA sequencing of the transcriptome revealed that *Nlgn3* transcripts are enriched in glial cell types including astrocytes and oligodendrocytes (Zhang et al., 2014). Interestingly, NLGNs expressed in astrocytes aid in neuron-astrocyte communication via bi-and tripartite synapses in mice in *C. elegans* (Hillen et al., 2018) suggesting that a neuronal-glial communication role may not be specific to mice. Similarly, NLGN3 expressed in enteric glia cloud also involve in establishing bi-partite and tripartite synapses in the ENS. Given this evidence, this study strongly suggests that NLGN3 might play a role in synapse formation and modifying synaptic function during neuron-glia communication in the ENS.

The impact of the R451C mutation on *Nlgn3* expression in glia has not previously been reported. Here we show that in *Nlgn3*^R451C^ mice, *Nlgn3* mRNA expression was reduced in myenteric glia but was unchanged in glia located within the submucosal plexus. Since glia play an important role in mediating intestinal functions such as mucosal barrier regulation, gut motility, immune resposes and neurotransmission such a reduction in NLGN3 levels could contribute to GI dysfunction in *Nlgn3*^R451C^ mutant mice. Therefore, in addition to neuronal dysfunction, glial dysfunction could also contribute to GI pathology in individuals diagnosed with ASD.

## CONCLUSION

This study aimed to characterise the effects of the *Nlgn3* R451C mutation on *Nlgn3* mRNA expression in the enteric nervous system. The dual RNAScope and immunofluorescence approach is a promising platform for characterising the expression of mRNA encoding synaptic proteins in the GI tract. We show that *Nlgn3* mRNA is expressed in most enteric neuronal and glial populations. In the myenteric plexus, mRNA expression is reduced in glia in *Nlgn3*^R451C^ mice, suggesting that the R451C mutation differentially modulates gene expression in the ENS. Taken together, these findings suggest that NLGN3 might play an important role in regulating ENS-mediated GI physiology and that changes to *Nlgn3* mRNA expression could contribute to GI dysfunction in individuals with ASD.

## Supplementary Information

**Figure S1.**
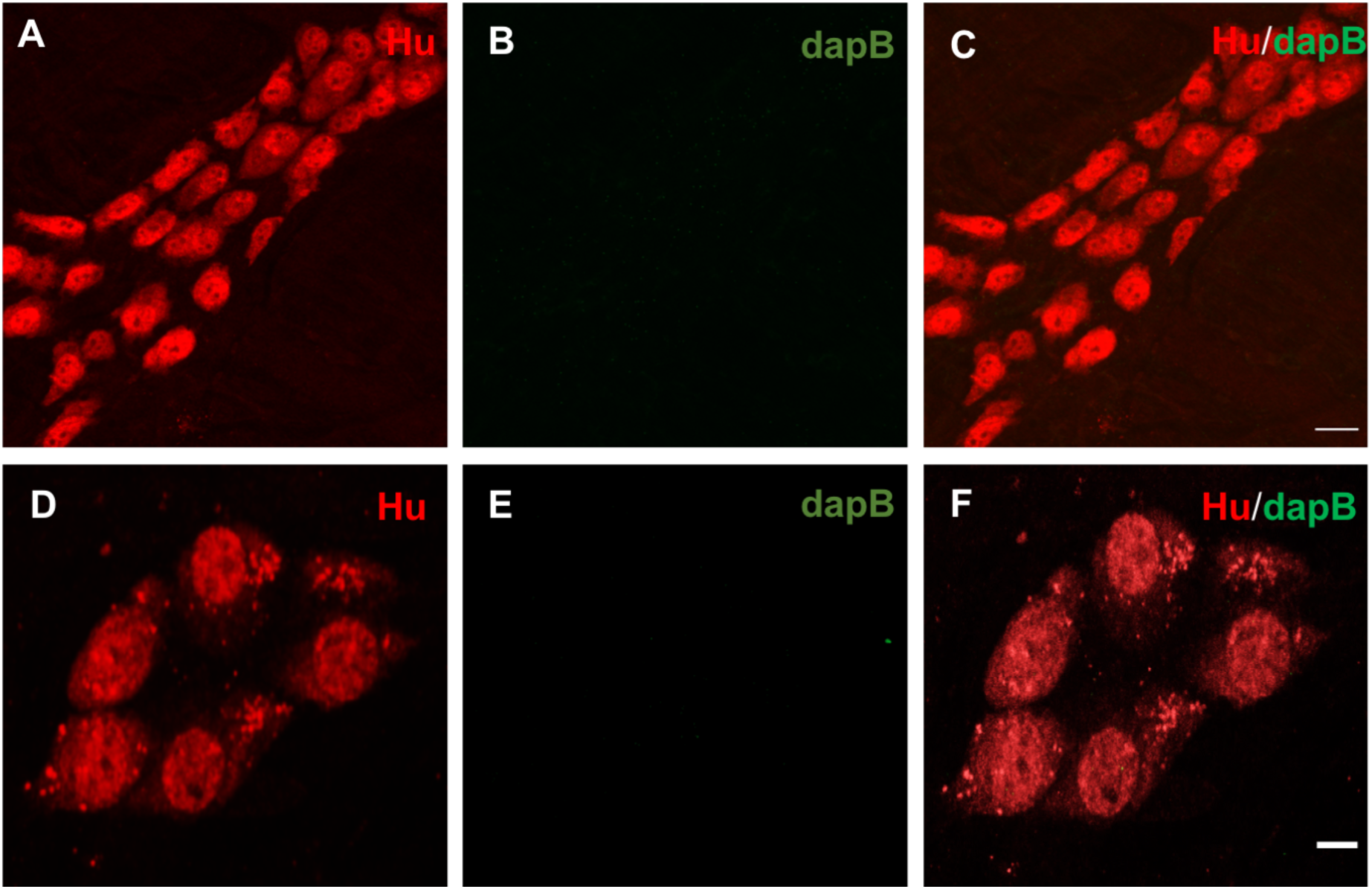
Negative control experiments for the myenteric plexus and the submucosal plexus. Myenteric neurons label with (A) Hu pan neuronal marker (B) negative control probe dapB. Double labelling of myenteric neurons with Hu and daPB. Submucosal neurons label with (D) Hu (B) dapB and co-labelling of myenteric neurons with Hu and dapB probe, scale bar=50µm.

**Figure S2.**
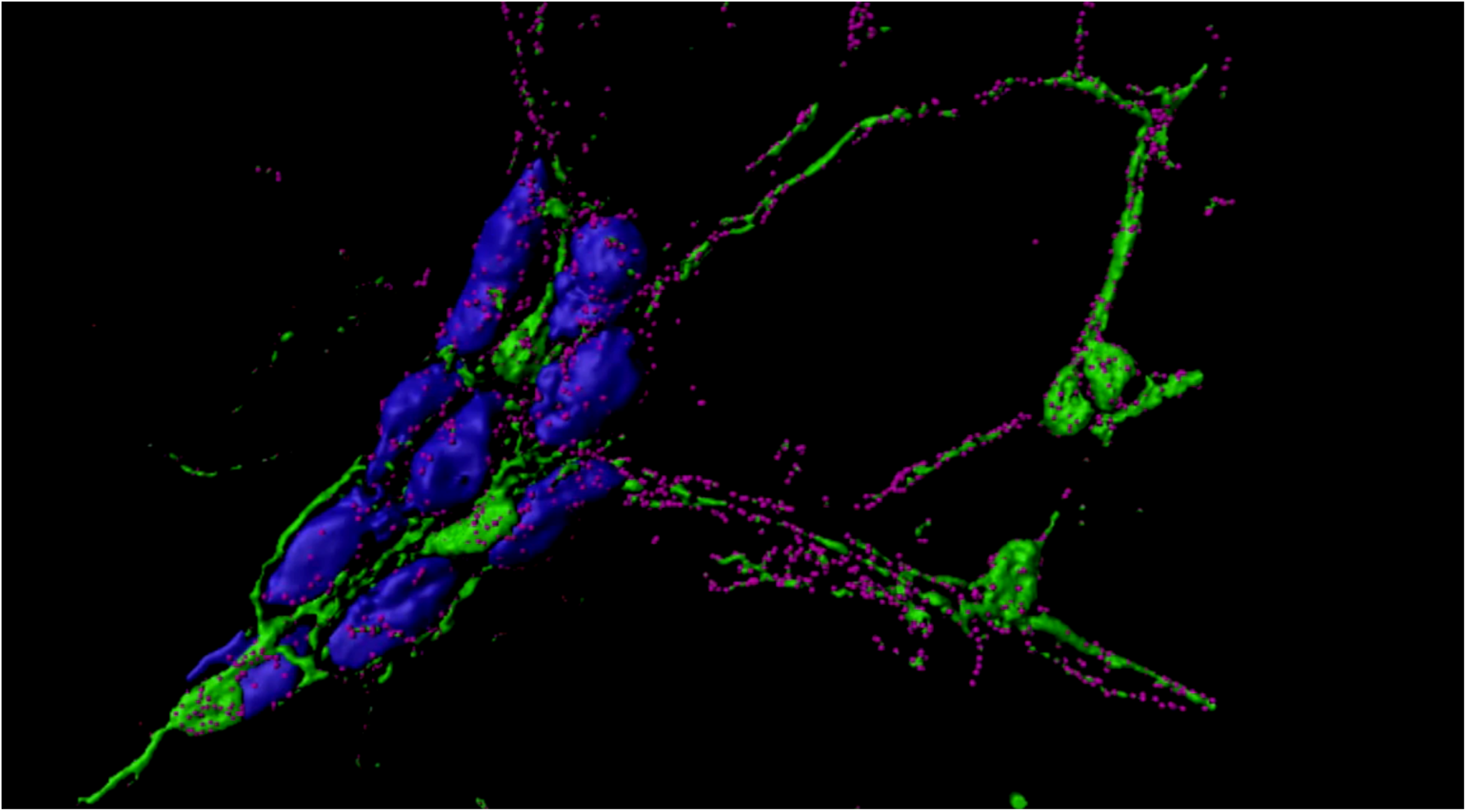
*Nlgn3* mRNA expression in submucosal neurons, glia and glial fibers. 3D reconstructed image of a mouse submucosal ganglion show *Nlgn3* mRNA expression in glial fibers.

